# Spindle assembly in egg extracts of the Marsabit clawed frog, *Xenopus* borealis

**DOI:** 10.1101/254979

**Authors:** Maiko Kitaoka, Rebecca Heald, Romain Gibeaux

## Abstract

Egg extracts of the African clawed frog *Xenopus laevis* have provided a cell-free system instrumental in elucidating events of the cell cycle, including mechanisms of spindle assembly. Comparison with extracts from the diploid Western clawed frog, *Xenopus tropicalis,* which is smaller at the organism, cellular and subcellular levels, has enabled the identification of spindle size scaling factors. We set out to characterize the Marsabit clawed frog, *Xenopus borealis,* which is intermediate in size between the two species, but more recently diverged in evolution from *X. laevis* than *X. tropicalis*. *X. borealis* eggs were slightly smaller than those of *X. laevis*, and slightly smaller spindles were assembled in egg extracts. Interestingly, microtubule distribution across the length of the *X. borealis* spindles differed from both *X. laevis* and *X. tropicalis*. Extract mixing experiments revealed common scaling phenomena among *Xenopus* species, while characterization of spindle factors katanin, TPX2, and Ran indicate that *X. borealis* spindles possess both *X. laevis* and *X. tropicalis* features. Thus, *X. borealis* egg extract provides a third *in vitro* system to investigate interspecies scaling and spindle morphometric variation.

## INTRODUCTION

In all eukaryotes, cell division requires the spindle, a highly dynamic, self-organizing, microtubule-based apparatus that accurately segregates replicated chromosomes to daughter cells. Spindle microtubules assemble into a bipolar structure that attaches to chromosomes via their kinetochores and aligns them at the metaphase plate so that sister chromatids are oriented toward opposite spindle poles. Although the basic form and composition of the spindle is conserved, spindle structure adapts to changes in cell size and type, and varies dramatically both across and within species (Wühr *et al.*, 2008; Crowder *et al.*, 2015). While many of the molecules and mechanisms that orchestrate spindle assembly have been elucidated, it remains unclear how specific spindle architectures are established, and how this variation impacts spindle function.

Unfertilized eggs of the African clawed frog *Xenopus laevis* arrested in metaphase of meiosis II by cytostatic factor (CSF) (Masui and Markert, 1971) can be obtained in large quantities. Their fractionation by centrifugation yields a crude cytoplasmic extract that maintains the metaphase arrest and supports spindle assembly around sperm nuclei or chromatin-coated beads (Sawin and Mitchison, 1991; Heald *et al.*, 1996). Remarkably, spindles formed in egg extracts of the smaller, related frog *Xenopus tropicalis* are shorter in length independent of DNA source, and mixing the two extracts produced spindles of intermediate size (Brown *et al.*, 2007). It was found that compared to *X. laevis*, *X. tropicalis* possess smaller spindles due to (i) elevated activity of the hexameric AAA-ATPase microtubule severing enzyme katanin (Loughlin *et al.*, 2010, 2011), and (ii) a higher concentration of the spindle assembly factor, TPX2 (targeting factor for Xklp2), which promotes association of the cross-linking spindle motor Eg5 and a shift of microtubule density from antiparallel overlap in the center of the spindle to the poles (Helmke and Heald, 2014). *X. tropicalis* spindles were also observed to possess distinct morphological features, including more robust kinetochore fibers and a more circular shape (Loughlin *et al.*, 2011; Grenfell *et al.*, 2016). Furthermore, whereas spindle assembly in *X. laevis* extracts required a chromatin-generated gradient of RanGTP (Kalab *et al.*, 2002; Cavazza and Vernos, 2016), *X. tropicalis* spindle assembly was not affected by disruption of this pathway (Helmke and Heald, 2014). Therefore, not only spindle size, but also spindle architecture, morphology, and assembly mechanisms differ between these two species.

To further address the conservation of spindle morphology, assembly, and scaling mechanisms across evolution, we decided to characterize a third *Xenopus* species, the Marsabit clawed frog *Xenopus borealis.* Whereas allotetraploid *X. laevis* (36 chromosomes) and diploid *X. tropicalis* (20 chromosomes) diverged 48 million years ago, *X. borealis* is more closely related to *X. laevis,* having diverged 17 million years ago (Session *et al.*, 2016). Like *X. laevis*, *X. borealis* possesses an allotetraploid genome of 36 chromosomes. Interestingly, *X. borealis* was used extensively in the 1970s to analyze the conservation of ribosomal nucleic acids compared to *X. laevis* (Brown *et al.*, 1972; Brown and Sugimoto, 1974; Griswold *et al.*, 1974; Leister and Dawid, 1975; Wellauer and Reeder, 1975; Ford and Brown, 1976) and was misidentified as *Xenopus mulleri* until 1977 (Brown *et al.*, 1977). Revisiting this species and applying the egg extract methodology allowed us to further evaluate spindle morphometric variation and scaling in *Xenopus*. We show that *X. borealis* spindles are intermediate in size and display properties of both *X. laevis* and *X. tropicalis*, resulting in distinct features. These findings highlight the evolutionary plasticity of spindle morphology and assembly mechanisms, and introduce *X. borealis* as a third *Xenopus* species amenable to *in vitro* assays to investigate the molecular basis of spindle variation.

## RESULTS AND DISCUSSION

In this study, we characterized *X. borealis* and spindle assembly in its egg extracts, using the two well-characterized *Xenopus* species, *X. laevis* and *X. tropicalis*, for comparison.

### *X. borealis* frogs, eggs, and sperm are morphologically distinct

At ∼7 cm body length, adult *X. borealis* adult frogs were intermediate in size between *X. laevis* (∼10 cm) and *X. tropicalis* (∼4 cm) and also displayed distinct physical traits including googly eyes and a pear-like body shape (**Figure 1A**). Measuring the size of *X. borealis* gametes revealed that *X. borealis* eggs averaged 1.2 ± 0.02 mm in diameter, slightly smaller than *X. laevis* (1.3 ± 0.05 mm) and larger than *X. tropicalis* (0.75 ± 0.02 mm) (**Figure 1B**). Interestingly, *X. borealis* sperm cells were significantly longer (average 22 ± 2.8 μm) than those of both *X. laevis* (19 ± 2.3 μm) and *X. tropicalis* (13.5 ± 2.1 μm) (**Figure 1C**), and similar size differences were observed comparing isolated sperm nuclei (**Supplementary Figure S1**) (Grainger, 2012). *In vitro* fertilization and live imaging revealed that the rate of early development of *X. borealis* and *X. laevis* embryos were similar (**Figure 1D and Video 1**) (Nieuwkoop and Faber, 1994). Nuclei isolated from stage 8 (5 hpf) embryos of *X. borealis* (average 2214 ± 1016 μm^2^) were comparable in size to those from *X. laevis* (2010 ± 1081 μm^2^), while *X. tropicalis* nuclei (405 ± 116 μm^2^) were significantly smaller (**Figure 1E**). In summary, although *X. borealis* display distinct traits, our measurements show that like *X. laevis* and *X. tropicalis*, *X. borealis* egg size scales with adult animal size, and that *X. borealis* development and nuclear size is very similar to that of *X. laevis*. These findings are consistent with the well-characterized phenomenon that genome size scales with nuclear and cell size among amphibians (Levy and Heald, 2016).

**FIGURE 1:**
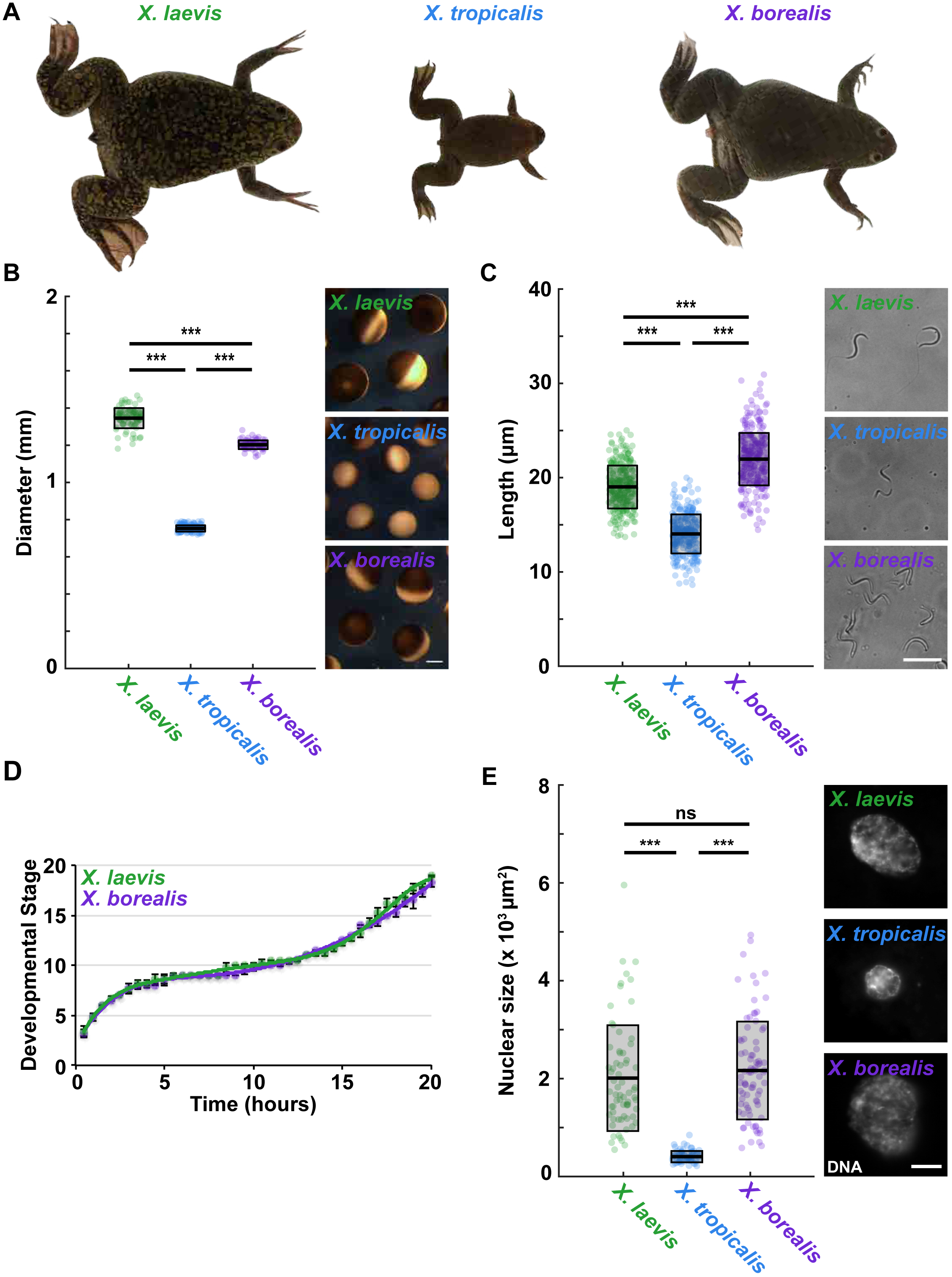
Characterization of the Marsabit clawed frog, *X. borealis*. **(A)** Relative body size of *X. borealis* frogs compared to *X. laevis* and *X. tropicalis*. **(B)** Comparison and quantification of egg size. Egg size (diameter in mm) is plotted as boxplots for *X. laevis* (green), *X. tropicalis* (blue), and *X. borealis* (purple). Each dot represents an individual egg (n = 89 for *X. laevis*, n = 132 for *X. tropicalis*, and n = 77 for *X. borealis*, from 3 females for each species). Representative images of eggs for each species are shown (right). Scale bar is 200 μm. **(C)** Comparison and quantification of sperm size. Sperm head length (μm) is plotted as boxplots for *X. laevis* (green), *X. tropicalis* (blue), and *X. borealis* (purple). Each dot represents an individual sperm cell (n = 288 for *X. laevis*, n = 278 for *X. tropicalis*, and n = 310 for *X. borealis*, from 3 males for each species). Representative images of sperm cells for each species are shown (right). Scale bar is 20 μm. **(D)** Early embryo development at room temperature for *X. laevis* and *X. borealis*. Embryos were staged according to Nieuwkoop and Faber (Nieuwkoop and Faber, 1994). Data for *X. laevis* embryos (n = 12) are blotted in green and *X. borealis* (n = 12) in purple. Error bars show the standard deviation. **(E)** Stage 8 embryo nuclei comparison and quantification. Representative images of nuclei isolated from stage 8 *X. laevis*, *X. tropicalis*, and *X. borealis* embryos are shown with DNA signal. Scale bar is 10 μm. Nuclear size (area in μm^2^) is plotted as boxplots for *X. laevis* (green), *X. tropicalis* (blue), and *X. borealis* (purple). Each dot represents an individual nucleus (n = 68 for *X. laevis*, n = 71 for *X. tropicalis*, and n = 84 for *X. borealis*). In **B**, **C**, and **E**, the thicker black line indicates the average, and the outlined gray box represents the standard deviation. Statistical significance was determined by a two-tailed, two-sample unequal variance t-test.

### *X. borealis* egg extract recapitulates cell cycle events

To evaluate the utility of *X. borealis* eggs for *in vitro* assays, we adapted published ovulation procedures to reproducibly obtain metaphase II, CSF-arrested eggs in sufficient amounts to prepare egg extracts (see **Material and Methods**). *X. borealis* eggs were dejellied and fractionated using standard protocols (Hannak and Heald, 2006; Maresca and Heald, 2006; Brown *et al.*, 2007), and produced similar yields of cytoplasmic extract compared to *X. laevis* despite their smaller body and egg size. *X. borealis* egg extract was then combined with *X. borealis* sperm nuclei and assayed for spindle assembly. When added directly to CSF extract, sperm nuclei induced the formation of asters (t = 5’), followed by half-spindles (t = 15’) and bipolar CSF spindles (t = 30 - 45’) (**Figure 2A, B**). Addition of calcium together with sperm nuclei induced exit from metaphase arrest and formation of interphase nuclei within 45 min. Following the addition of fresh CSF extract, bipolar “cycled” spindles assembled from 75 min to 90 min (**Figure 2A, C**). We noted that *X. borealis* extracts formed spindles more quickly than *X. laevis*, almost as rapidly as *X. tropicalis* (unpublished data). These experiments show that, as for the other *Xenopus* species, *X. borealis* egg extracts are fully capable of recapitulating the cell cycle and assembling spindles *in vitro.*

**FIGURE 2:**
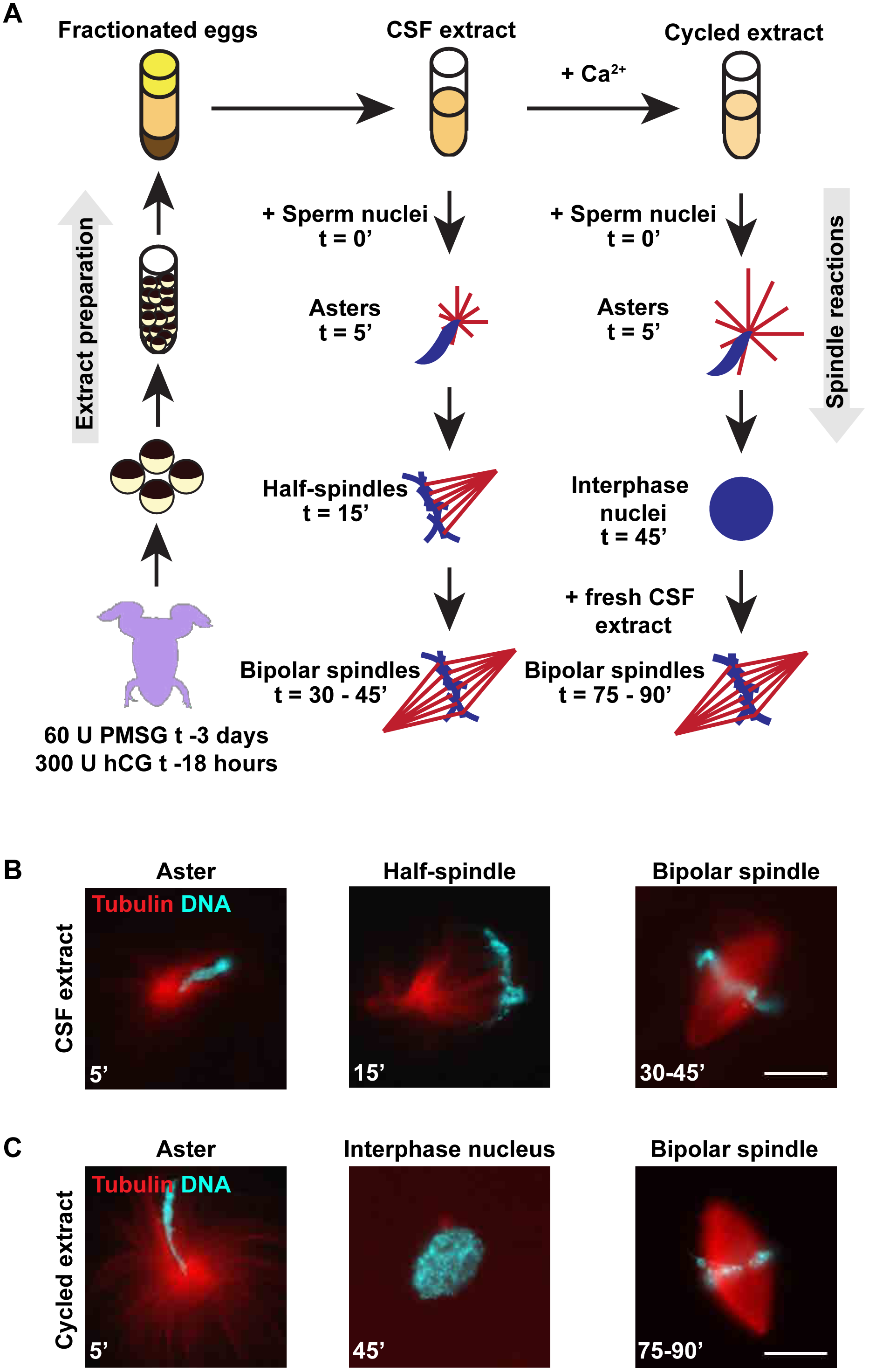
*X. borealis* egg extracts at a glance. **(A)** Schematic of extract preparation and spindle assembly reactions. **(B)** Images of spindle assembly intermediates in CSF extract reactions. **(C)** Images of spindle assembly intermediates in cycled extract reactions. Images in **B-C** of representative structures are shown with tubulin (red) and DNA (blue) signal. Scale bar is 20 μm.

### *Xenopus* species possess distinct spindle morphometrics

We next examined some parameters of *X. borealis* spindle size and microtubule organization and compared them to *in vivo* meiosis II spindles as well as to spindles assembled in *X. laevis* and *X. tropicalis* egg extracts. *X. borealis* spindles assembled *in vitro* were similar in length from pole-to-pole and width at the metaphase plate to meiotic spindles imaged in metaphase II-arrested eggs (length/width average 43.9 ± 5.5 μm/21.7 ± 3.6 μm *in vivo* vs. 44.4 ± 5.5 μm/23.4 ± 5.9 μm in extract) (**Figure 3A**). *X. borealis* extract was also capable of inducing bipolar spindle assembly around stage 8 *X. borealis* embryo nuclei and 10 μm diameter chromatin-coated beads (Halpin *et al.*, 2011). Embryo nuclei spindles were slightly smaller than those formed around sperm nuclei (average length 39.7 ± 4.4 μm, width 25.1 ± 4.7 μm) (**Figure 3B**), while spindles assembled around DNA-coated beads were considerably shorter (average 22.4 ± 3.9 μm) (**Figure 3C**). These measurements demonstrate that like *X. laevis* and *X. tropicalis*, spindles formed in *X. borealis* egg extract mimic the size and morphology of the meiotic egg spindle, and that spindle assembly *in vitro* occurs in the presence or absence of centrosomes.

**FIGURE 3:**
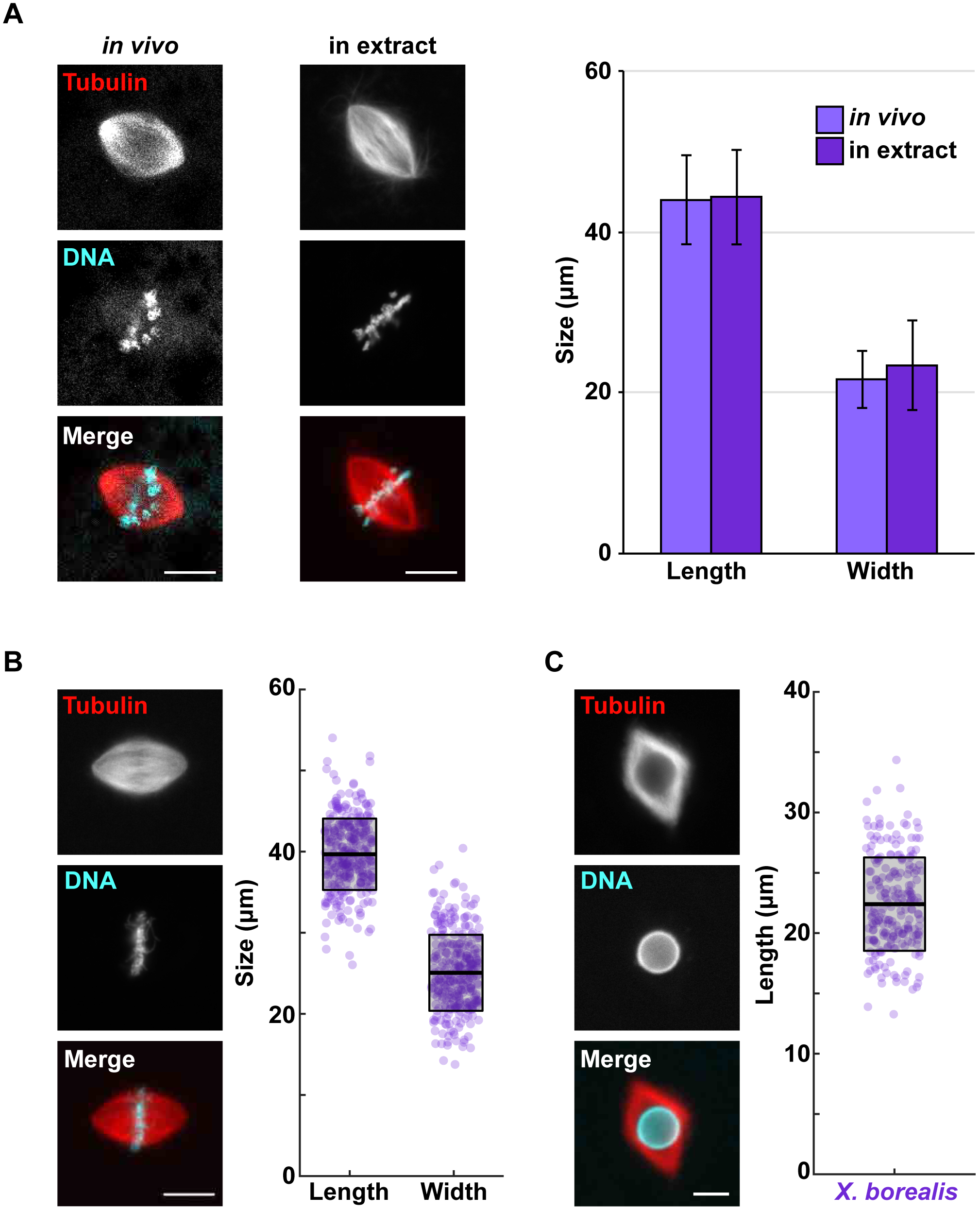
*X. borealis* CSF egg extracts reconstitute spindle assembly. **(A)** *In vivo X. borealis* metaphase-arrested egg spindles compared to spindles assembled in CSF extract. Scale bar is 20 μm. Spindle length (left) and width (right) are plotted (μm) as columns for *X. borealis* spindles from *in vivo* metaphase-arrested eggs (light purple) and formed in *in vitro* egg extract (dark purple). Error bars represent the standard deviation. **(B)** Spindles assembled around stage 8 *X. borealis* embryo nuclei added to *X. borealis* egg extract. Scale bar is 20 μm. Spindle length (left) and width (right) are plotted (μm) as boxplots. Each dot represents an individual spindle measurement (n = 310, from 3 independent egg extracts). **(C)** Spindles assembled around single 10 μm chromatin-coated beads in *X. borealis* egg extract. Scale bar is 10 μm. Length (μm) is plotted as a boxplot. Each dot represents an individual spindle measurement (n = 219, from 4 independent egg extracts). Representative images in **A-C** are shown with tubulin (top) and DNA (middle) signals individually and merged (bottom). In **B-C**, the thicker black line indicates the average, and the outlined gray box represents the standard deviation.

Despite their smaller egg size, we found that cycled *X. borealis* spindles were similar in length to *X. laevis* (average 41.9 ± 4.1 μm vs. 41.6 ± 6.0 μm), but significantly narrower (average 19.7 ± 3.7 μm vs. 22.4 ± 3.6 μm), thereby reducing spindle area and therefore total microtubule content. *X. borealis* spindles were both longer and wider than *X. tropicalis* spindles (average 23.2 ± 3.0 μm in length, 10.4 ± 2.6 μm in width) (**Figure 4A**). Thus, *X. borealis* spindles scale with egg size and are intermediate in size between those of *X. laevis* and *X. tropicalis*. Interestingly, the three species each displayed unique spindle microtubule distributions across the spindle. As previously reported, *X. laevis* spindles showed a consistent plateau of tubulin intensity along the length of the spindle indicating the presence of an overlapping, tiled array of microtubules (Loughlin *et al.*, 2010). In contrast, *X. tropicalis* tubulin intensity was greater at the poles and reduced in the center of the spindle at the metaphase plate (Helmke and Heald, 2014). Interestingly, *X. borealis* spindles displayed a similar increase in microtubule density at the poles as *X. tropicalis*, a similar density at the metaphase plate as *X. laevis*, and a unique dip in tubulin intensity between the poles and metaphase plate (**Figure 4B**). While similar microtubule distribution patterns were observed in CSF spindles, they were less distinct than in cycled spindles, potentially due to the absence of sister kinetochores and associated microtubules (Grenfell *et al.*, 2016). *X. borealis* CSF spindles were also longer and wider than both *X. laevis* and *X. tropicalis* (**Supplementary Figure S2**). A difference in spindle morphology between spindles with replicated (cycled) and unreplicated (CSF) chromosomes has been observed previously and emphasizes the fact that spindle morphology can vary in different cellular contexts (Grenfell *et al.*, 2016; Levy and Heald, 2016). We decided to focus on cycled spindle assembly for subsequent experiments, as these form around duplicated chromosomes with assembled kinetochores and thus are more physiologically relevant. Therefore, *X. borealis* fits into the *Xenopus* interspecies scaling regime, as the animals, eggs and meiotic spindles are all intermediate in size between the other two species. However, *X. borealis* microtubule arrays share some similarity with both species, leading to a unique spindle architecture.

**FIGURE 4:**
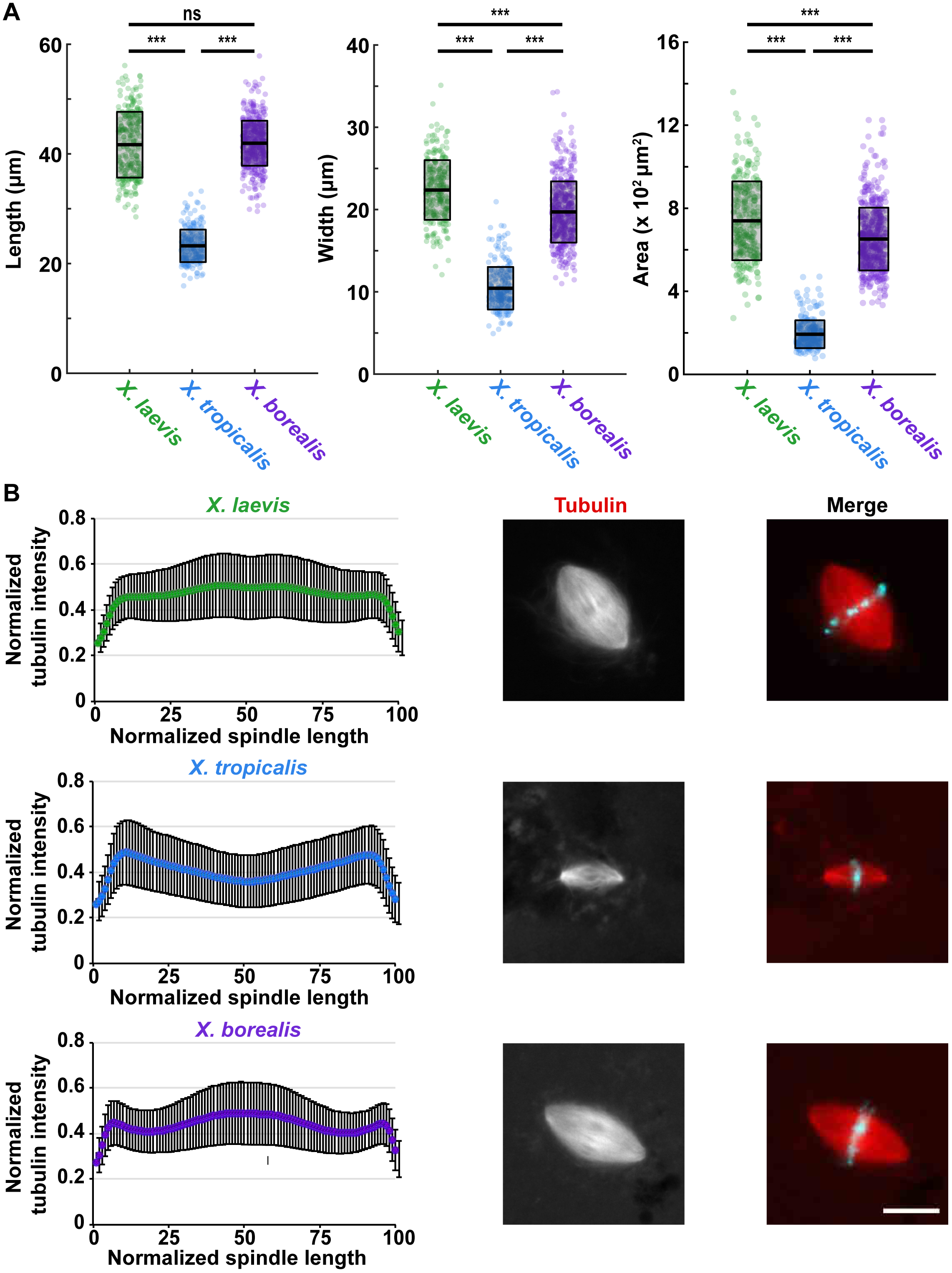
Analysis of *X. borealis* cycled spindles. **(A)** Quantification of *X. borealis* cycled spindle dimensions. Length (left), width (middle), and area approximated as an ellipse (right) are plotted (μm or μm^2^) as boxplots for *X. laevis* (green), *X. tropicalis* (blue), and *X. borealis* (purple) spindles. Each dot represents an individual spindle measurement (n = 256 spindles for *X. laevis*, n = 241 for *X. tropicalis*, and n = 493 for *X. borealis*, from 3 independent egg extracts for each species), the thicker black line indicates the average, and the outlined gray box represents the standard deviation. Statistical significance was determined by a two-tailed, two-sample unequal variance t-test. **(B)** Microtubule fluorescence intensity distribution in *X. laevis* (green), *X. tropicalis* (blue), and *X. borealis* (purple) spindles. Line scans of rhodamine-tubulin signal along the length of the spindle were measured (n = 257 spindles for *X. laevis*, n = 242 for *X. tropicalis*, and n = 470 for *X. borealis*, from 3 independent egg extracts for each species). Spindle length was normalized to 100% and tubulin intensities were normalized within each dataset. Average tubulin intensities were plotted for each species’ spindles. Error bars represent the standard deviation. Representative images of spindles formed from *X. laevis, X. tropicalis*, and *X. borealis* sperm nuclei in their respective egg extracts are shown with tubulin signal individually (left) and merged with DNA signal (right). Scale bar is 20 μm.

### *X. borealis* spindles are sensitive to perturbation of the RanGTP gradient

To compare spindle assembly mechanisms among the three species, we examined the role of the RanGTP gradient, which acts to release nuclear localization signal (NLS)-containing spindle assembly factors from importins near mitotic chromatin where the guanine nucleotide exchange factor RCC1 (Regulator of Chromosome Condensation 1) is localized, thereby promoting microtubule nucleation, stabilization, and crosslinking (Cavazza and Vernos, 2016). Perturbing the RanGTP gradient by the addition of Ran mutants, such as a dominant negative (T24N) or constitutively active, GTP-bound (Q69L) form, is detrimental to *X. laevis* spindle assembly, abolishing it completely or increasing ectopic microtubule nucleation and formation of multipolar spindles, respectively (Kalab *et al.*, 1999, 2002). In contrast, we showed previously that *X. tropicalis* spindles are much less affected by perturbation of the RanGTP gradient (Helmke and Heald, 2014). To test the role of this pathway in *X. borealis*, we added the Ran mutants at the onset of spindle assembly. Upon addition of RanT24N, spindle assembly was strongly impaired, with dramatically reduced microtubule nucleation around sperm nuclei, while RanQ69L increased the number of multipolar spindle structures (**Figure 5 and Supplementary Table 1)**. Thus, like *X. laevis*, *X. borealis* bipolar spindle assembly depends on the RanGTP gradient.

**FIGURE 5:**
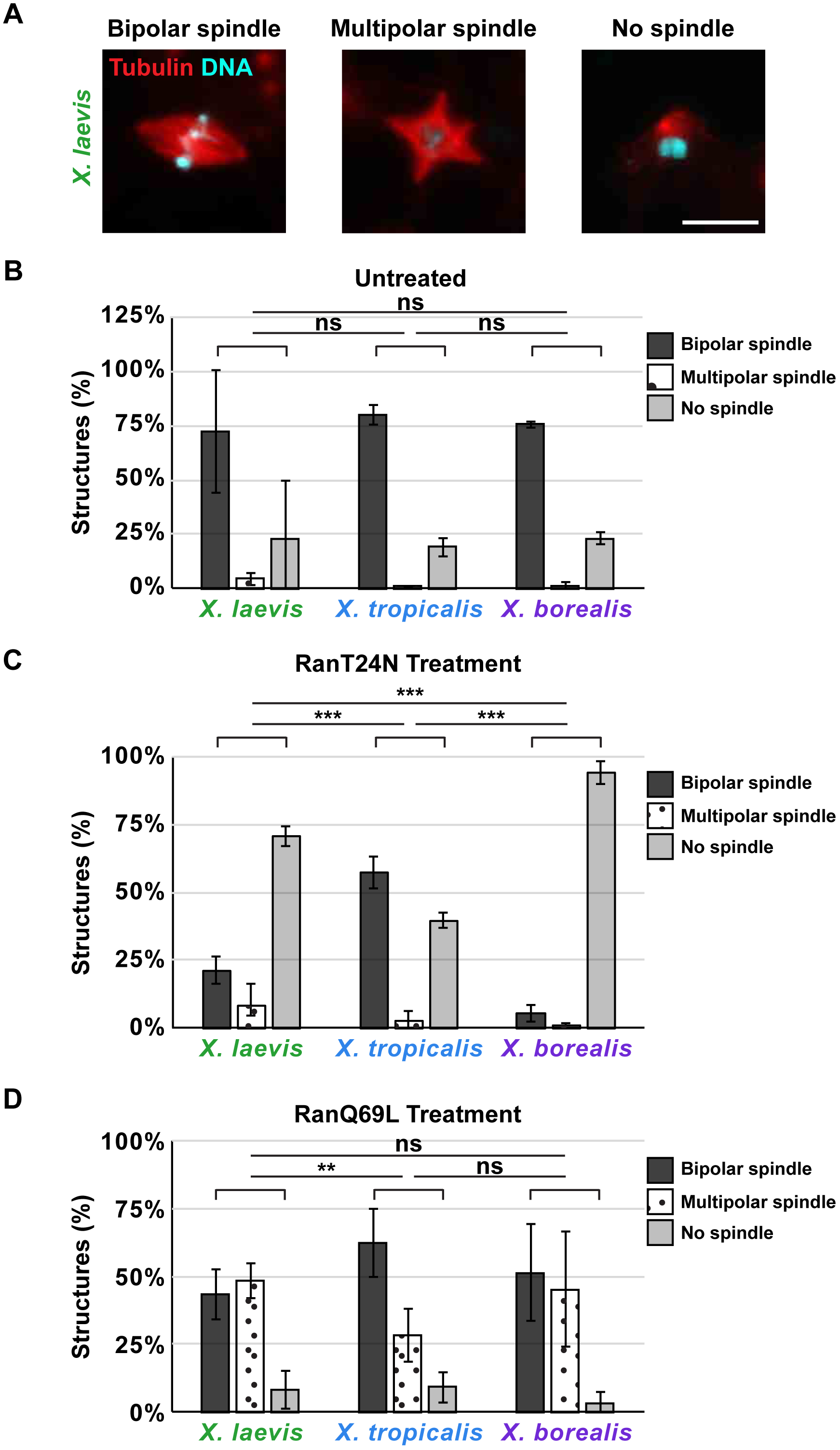
Sensitivity of *X. borealis* spindles to perturbation of the RanGTP gradient. **(A)** Phenotype categories quantified in *Xenopus* egg extract spindle assembly reactions containing Ran mutants. Representative images in *X. laevis* extracts are shown with tubulin (red) and DNA (blue) signals. Scale bar is 20 μm. Bar graphs of the percentage of spindles in each category are shown for **(B)** untreated extracts, **(C)** RanT24N treated extracts, and **(D)** RanQ69L treated in *Xenopus* extracts. In **B-D**, percentages of bipolar (dark gray), multipolar (polka dots), and no spindles found (light gray) are plotted for *X. laevis*, *X. tropicalis*, and *X. borealis* extracts (3 independent extracts were quantified for each species). Error bars represent the standard deviation. Statistical significance was determined by a two-tailed, 3x2 Fisher Exact test.

### Common spindle scaling mechanisms operate among *Xenopus* species

It has been shown that hybrid embryos can be generated by fertilization of *X. borealis* eggs with sperm from either *X. laevis* and *X. tropicalis* (De Robertis and Black, 1979; Gibeaux *et al.*, 2018), indicating compatibility between the *X. borealis* cytoplasm and the chromosomes of both species. Addition of either *X. laevis* or *X. tropicalis* sperm nuclei to *X. borealis* extract did not alter spindle microtubule distribution (**Supplementary Figure S3A)**, consistent with previous studies indicating that egg cytoplasm composition is the primary determinant of spindle morphology (Brown *et al.*, 2007). To determine whether maternal cytoplasmic factors affect spindle size similarly among the three *Xenopus* species, we mixed *X. borealis* egg extract with either *X. laevis* or *X. tropicalis* extract in the presence of *X. laevis* or *X. tropicalis* sperm nuclei, respectively. Whereas titration of *X. borealis* extract with *X. laevis* did not affect spindle dimensions (**Figure 6A**), *X. tropicalis* extract caused *X. borealis* spindles to shrink proportionately with the amount added (**Figure 6B**). Analysis of spindle microtubule distributions in mixed extracts revealed that these were also affected proportionately, favoring the tubulin distribution of the more abundant extract (**Supplementary Figure S3B and C**). These results are consistent with cytoplasmic mixing of *X. laevis* with *X. tropicalis* extracts, which also caused spindle shrinkage in a dose-dependent manner (Brown *et al.*, 2007). Altogether, these data indicate that common scaling and microtubule organization mechanisms operate among the three *Xenopus* species.

**FIGURE 6:**
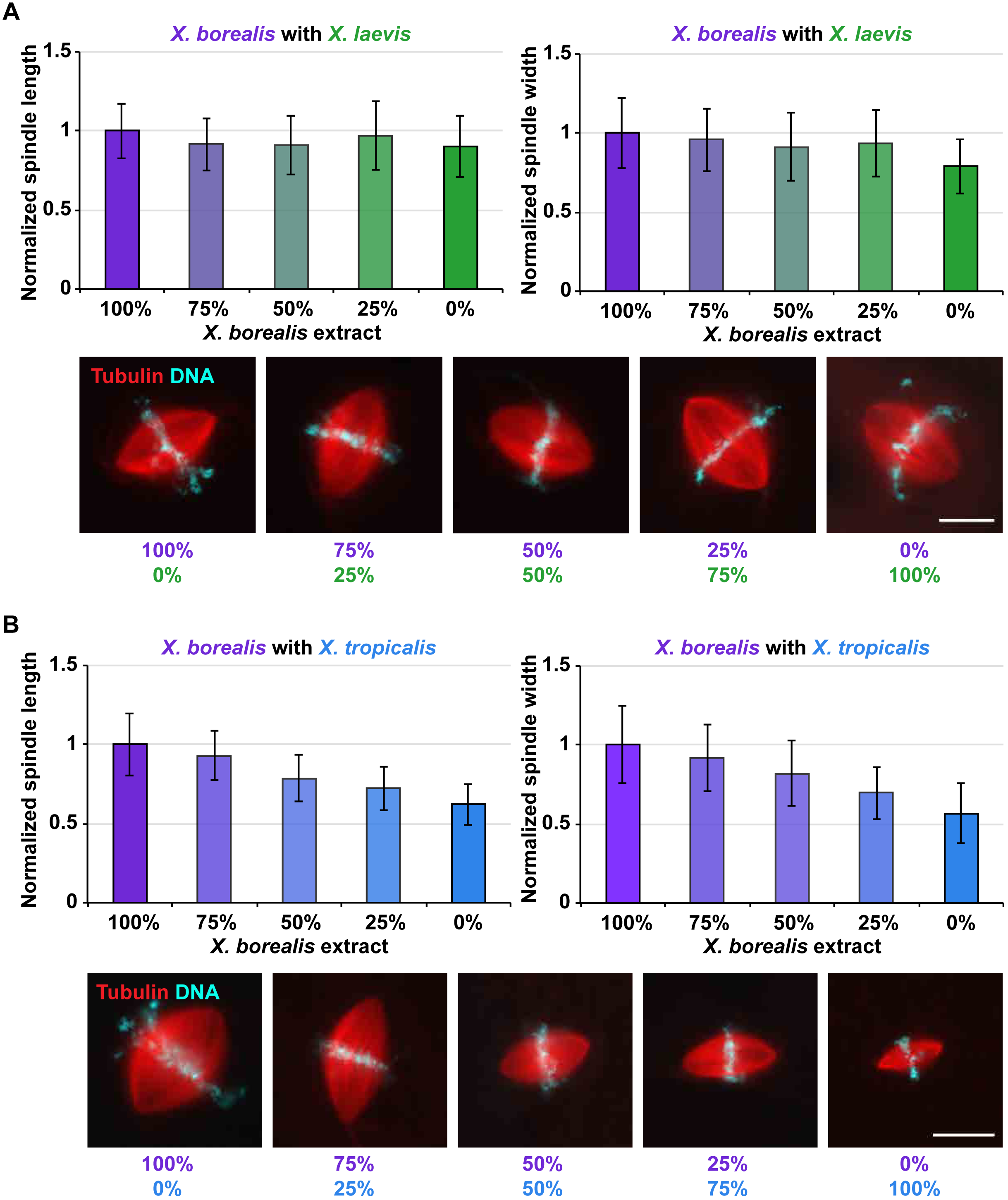
Titration of *X. borealis* egg extract with other *Xenopus* egg extracts. **(A)** *X. borealis* extract mixed with *X. laevis* extract. Spindle length (left panel) and width (right panel) are plotted as columns, with colors proportionate to the amount of *X. laevis* (green) added to *X. borealis* (purple) (n = 341 for 100% *X. borealis*, 0% *X. laevis*; n = 350 for 75% *X. borealis*, 25% *X. laevis*; n = 331 for 50% *X. borealis*, 50% *X. laevis*; n = 313 for 25% *X. borealis*, 75% *X. laevis*; and n = 267 for 0% *X. borealis*, 100% *X. laevis* over 3 independent experiments). **(B**) *X. borealis* extract mixed with *X. tropicalis* extract. Spindle length (left panel) and width (right panel) are plotted as columns, with colors proportionate to the amount of *X. tropicalis* (blue) added to *X. borealis* (purple) (n = 295 for 100% *X. borealis*, 0% *X. tropicalis*; n = 306 for 75% *X. borealis*, 25% *X. tropicalis*; n = 295 for 50% *X. borealis*, 50% *X. tropicalis*; n = 337 for 25% *X. borealis*, 75% *X. tropicalis*; and n = 334 for 0% *X. borealis*, 100% *X. tropicalis* over 3 independent experiments). In **A-B**, spindle length and width were normalized to the length and width, respectively, of 100% *X. borealis* CSF spindles. Error bars represent the standard deviation. Representative images are shown with tubulin (red) and DNA (blue) signal. Scale bar is 20 μm.

### *X. borealis* spindles combine scaling factor features of both *X. laevis* and *X. tropicalis*

To investigate the molecular basis of *X. borealis* spindle size and morphology, we compared the localization, abundance, and sequence of katanin and TPX2, two spindle scaling factors that contribute to spindle size differences between *X. laevis* and *X. tropicalis*, as well as the microtubule polymerase XMAP215, levels of which have been shown to regulate spindle length in *Xenopus* (Reber *et al.*, 2013). XMAP215 levels and localization were similar for all three *Xenopus* species (**Supplementary Figure S4**). Katanin, a microtubule severing AAA-ATPase, has higher activity in *X. tropicalis* due to the absence of an inhibitory phosphorylation site (Serine 131) found in the *X. laevis* protein (Loughlin *et al.*, 2011). Immunofluorescence and line scan analysis of spindles revealed that katanin localization on the *X. borealis* spindle displays a small but distinct increase at the metaphase plate as well as the spindle poles, while the localization on *X. laevis* spindles is more uniform and *X. tropicalis* shows katanin enrichment solely at the poles (**Figure 7A**). By Western blot, katanin levels did not differ significantly across the three species (**Figure 7B**). Although the amino acid sequence of *X. borealis* katanin is over 97% identical to that of both *X. tropicalis* and *X. laevis*, it contains the key inhibitory serine residue that regulates katanin activity (**Supplementary Figure S5**). TPX2 is a Ran-regulated microtubule-associated protein that is found at three-fold higher concentrations in *X. tropicalis* compared to *X. laevis*. Increasing the level of TPX2 in *X. laevis* extracts to that of *X. tropicalis* was previously shown to reduce spindle length through its increased recruitment of the spindle motor Eg5 to spindle poles (Helmke and Heald, 2014). Interestingly, *X. borealis* spindles displayed an overall TPX2 localization more similar to *X. tropicalis*, with higher intensity at the spindle poles, but also possessed a central smaller peak of TPX2 reminiscent of *X. laevis* (**Figure 7C**). Levels of TPX2 in *X. borealis* egg extracts were two-fold higher than in *X. laevis,* but not elevated to *X. tropicalis* levels. (**Figure 7D**). Notably, the amino acid sequence of *X. borealis* TPX2 is more similar to *X. laevis* compared to *X. tropicalis* and contains the seven amino acid residues thought to reduce microtubule nucleation activity of *X. laevis* TPX2 (**Supplementary Figure S6**) (Helmke and Heald, 2014; Grenfell *et al.*, 2016). Altogether, these results suggest that *X. borealis* spindle length results from its lower katanin activity similar to *X. laevis*, while its microtubule distribution is likely affected by increased levels of TPX2, which is more reminiscent of *X. tropicalis*.

**FIGURE 7:**
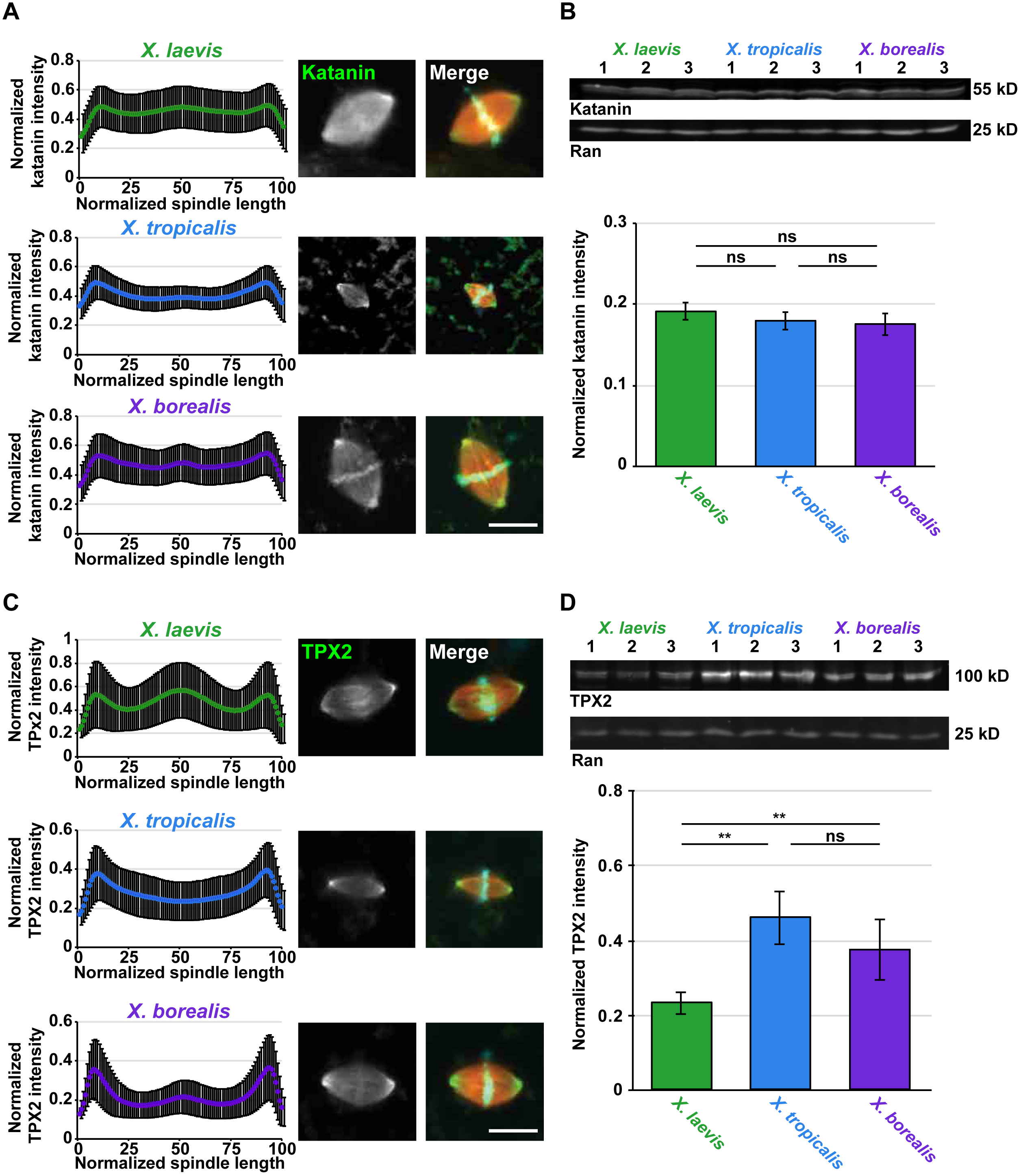
Localization of scaling factors katanin and TPX2 on *X. borealis* spindles. **(A)** Katanin immunofluorescence staining of *X. laevis* (green), *X. tropicalis* (blue), and *X. borealis* (purple) spindles. Line scans of Alexa Fluor 488 signal along the length of the spindle were measured (n = 385 spindles for *X. laevis*, n = 408 for *X. tropicalis*, and n = 348 for *X. borealis*, from 3 independent egg extracts for each species). **(B)** Western blots of *X. laevis*, *X. tropicalis*, and *X. borealis* extracts probed for katanin. **(C)** TPX2 immunofluorescence staining of *X. laevis* (green), *X. tropicalis* (blue), and *X. borealis* (purple) spindles. Line scans of Alexa Fluor 488 signal along the length of the spindle were measured (n = 406 spindles for *X. laevis*, n = 433 for *X. tropicalis*, and n = 418 for *X. borealis*, from 3 independent egg extracts for each species). **(D)** Western blots of *X. laevis*, *X. tropicalis*, and *X. borealis* extracts, probed for TPX2. In **A** and **C**, spindle length was normalized to 100% and katanin or TPX2 intensities, respectively, were normalized within each dataset. Average TPX2 intensities were plotted for each species’ spindles. Error bars represent the standard deviation. Representative images of spindles formed from *X. laevis, X. tropicalis*, and *X. borealis* sperm nuclei in their respective egg extracts are shown with katanin or TPX2 grayscale signal (left) and merged with tubulin and DNA signals (right). Scale bar is 20 μm. In **B** and **D**, katanin or TPX2 band integrated densities are plotted as columns for *X. laevis* (green), *X. tropicalis* (blue), and *X. borealis* (purple) extracts (3 independent extracts are shown for each species). Band intensities were normalized to the integrated density of the corresponding Ran loading control. Error bars represent the standard deviation. Statistical significance was determined by a two-tailed, two-sample unequal variance t-test.

Overall, these observations reveal that *X. borealis* meiotic spindles, although more similar in size and RanGTP-dependence to *X. laevis*, also possess some structural and molecular features characteristic of *X. tropicalis*. These findings support the idea that katanin and TPX2 are important spindle scaling and assembly factors in *Xenopus*. Katanin-mediated microtubule severing activity has been shown to regulate spindle size in several organisms, including *Caenorhabditis elegans*, *Xenopus*, and human cells (McNally *et al.*, 2006; Loughlin *et al.*, 2011; Joly *et al.*, 2016; Jiang *et al.*, 2017), which is consistent with results from meiotic spindle assembly simulations showing that altering microtubule depolymerization rates robustly scales spindle length (Loughlin *et al.*, 2010).

The role of TPX2 in spindle assembly and size control is more complicated. Depletion of the TPX2 ortholog D-TPX2 from *Drosophila melanogaster* syncytial embryos and S2 cells resulted in spindle shortening (Goshima, 2011; Hayward and Wakefield, 2014), and human cells require phosphorylation of TPX2 by Aurora A for normal spindle length (Fu *et al.*, 2015). In contrast, addition of recombinant TPX2 to *X. laevis* extracts decreased spindle length (Helmke and Heald, 2014), and we have observed similar effects on *X. borealis* spindles (unpublished data). Spindle-shrinking effects in *Xenopus* did not depend on Aurora A, but rather on a change in spindle microtubule distribution caused by increased recruitment of the cross-linking Eg5 motor to spindle poles (Helmke and Heald, 2014). TPX2 also regulates microtubule nucleation in the spindle (Petry *et al.*, 2013; Scrofani *et al.*, 2015; Petry, 2016; Alfaro-Aco *et al.*, 2017). TPX2 has thus emerged as a central player of spindle assembly and organization in a wide variety of cell types (Ma *et al.*, 2011; Helmke *et al.*, 2013; Fu *et al.*, 2015; Levy and Heald, 2016) and is overexpressed in many cancers (Chang *et al.*, 2012; Neumayer *et al.*, 2014). Our results highlight the importance of TPX2 levels and localization as determinants of meiotic spindle microtubule architecture. TPX2 localization (**Figure 7C**) correlates well with the distribution of tubulin intensity across all three species (**Figure 4B**), particularly at the spindle poles (Ma *et al.*, 2011; Helmke and Heald, 2014). Despite having higher TPX2 levels, the greater sequence similarity of *X. borealis* TPX2 to the *X. laevis* protein (**Supplementary Figure S6**) may explain the increased sensitivity to Ran inhibition in *X. borealis* (**Figure 5C**), since a 7 amino acid sequence shown to decrease microtubule nucleation activity relative to *X. tropicalis* is present in both *X. borealis* and *X. laevis* TPX2 (**Figure 7D**) (Helmke and Heald, 2014). Altogether, our results provide clues to how TPX2 levels may tune meiotic spindle architecture and assembly pathways by altering microtubule nucleation and organization.

In summary, *Xenopus borealis* provides a robust *in vitro* egg extract that adds to the characterized *X. laevis* and *X. tropicalis* systems, enabling further investigation of spindle assembly and interspecies scaling. Our findings highlight that egg meiosis II spindle architecture varies across *Xenopus* species and suggest that *X. borealis* spindles assemble with a combination of *X. laevis* as well as *X. tropicalis* features.

## MATERIALS AND METHODS

### Chemicals

Unless otherwise stated, all chemicals were purchased from Sigma-Aldrich, St. Louis, MO.

### Frog care

All animal experimentation is this study was performed according to our Animal Use Protocol approved by the UC Berkeley Animal Care and Use Committee. Mature *X. laevis*, *X. tropicalis*, and *X. borealis* frogs were obtained from NASCO, WI. *X. laevis*, *X. tropicalis*, and *X. borealis* females were ovulated with no harm to the animals with a 6-, 3-, and 4-month rest interval, respectively. To obtain testes, males were euthanized by over-anesthesia through immersion in ddH_2_O containing 0.15% MS222 (Tricaine) neutralized with 5 mM sodium bicarbonate prior to dissection, and then frozen at -20°C.

### Sperm nuclei purification

*X. laevis* and *X. tropicalis* sperm nuclei were prepared and purified as previously described (Grainger, 2012). The following modifications were made for *X. borealis* sperm nuclei purification. *X. borealis* males were primed with 40 U of PMSG 3 days prior to dissection and boosted with 100 U hCG 12-24 h before dissection. Testes were removed and cleaned from all blood vessels and fat with forceps. Testes were then washed 3X in cold 1X MMR (1X MMR: 100 mM NaCl, 2 mM KCl, 2 mM CaCl_2_, 1 mM MgCl_2_ and 5 mM HEPES-NaOH pH 7.6, 0.1 mM EDTA) and 3X in cold Nuclear Preparation Buffer (NPB) (2X NPB: 500 mM sucrose, 30 mM HEPES, 1 mM spermidine trihydrochloride, 0.4 mM spermine tetrahydrochloride, 2 mM dithiothreitol, 2 mM EDTA) and placed in 1 mL of cold NPB for homogenization with scissors and a pestle. Samples were briefly spun to pellet the remaining tissue before transferring the supernatant to a new tube for centrifugation at 1500 g for 10 min at 4°C to pellet sperm. The pellet was resuspended in 1 mL NPB supplemented with 50 μL of 10 mg/mL lysolecithin and incubated for 5 min at RT. The resuspended pellet was then added to 4 mL of cold NPB supplemented with 3% BSA and spun at 3,000 RPM for 10 min at 4°C. The resulting pellet was resuspended in 2 mL of cold NPB with 0.3% bovine serum albumin (BSA) and 30% filtered glycerol. Finally, the pellet was resuspended in 500 μL of cold NPB with 0.3% BSA and 30% filtered glycerol. Sperm nuclei were adjusted to a concentration of 10^8^ nuclei per mL using cold NPB with 0.3% BSA and 30% glycerol to make a 200X stock, aliquoted, frozen in liquid nitrogen, and stored at -80°C.

### *In vitro* fertilization

*X. borealis* males were injected with 300 U of human chorionic gonadotropin hormone (hCG) 12-24 h before dissection and testes were collected in Leibovitz L-15 Medium (Gibco – Thermo Fisher Scientific, Waltham, MA) supplemented with 10% Fetal Bovine Serum (FBS; Gibco) for immediate use. *X. laevis* males were injected with 500 U of hCG 12-24 h before dissection and testes were stored at 4°C in 1X MR (100 mM NaCl, 1.8 mM KCl, 2 mM CaCl_2_, 1 mM MgCl_2_ and 5 mM HEPES-NaOH pH 7.6) for 1-2 weeks.

*X. borealis* and *X. laevis* females were primed with 60 U and 100 U, respectively, of pregnant mare serum gonadotrophin (PMSG, National Hormone and Peptide Program, Torrance, CA) at least 48 h before use and boosted with 300 U and 500 U, respectively, of hCG 12-24 h before the experiment. *X. borealis* and *X. laevis* frogs were kept at 16°C in 0.5X and 1X MMR, respectively. *X. borealis* eggs were picked from the tub and deposited onto petri dishes coated with 1.5% agarose in 1/10X MMR. *X. laevis* females were squeezed gently to deposit eggs onto petri dishes coated with 1.5% agarose in 1/10X MMR. Two *X. borealis* testes or 2/3 of an *X. laevis* testis were collected and homogenized using scissors and a pestle in 1 mL of L-15 and 10% FBS. Any excess liquid in the petri dishes was removed, and the eggs were fertilized with 500 μL of sperm solution per dish. Eggs were swirled in the solution to individualize eggs as much as possible and incubated for 5 min. Dishes were flooded with ddH_2_O, swirled and incubated for 10 min, followed by buffer exchange to 1/10X MMR and incubation for 10 min. The embryo jelly coats were then removed with a 3% cysteine solution in ddH_2_O-NaOH, pH 7.8. After extensive washing (>4X) with 1/10X MMR, embryos were incubated at 23°C. At stage 2-3, fertilized embryos were sorted and placed in fresh 1/10X MMR in new petri dishes coated with 1.5% agarose in 1/10X MMR.

All embryos were staged according to Nieuwkoop and Faber (Nieuwkoop and Faber, 1994).

### Embryo video imaging

Imaging dishes were prepared using a homemade PDMS mold designed to print a pattern of 1 mm large wells in agarose that allowed us to image 4 *X. borealis* or *X. laevis* embryos simultaneously within the 3X4 mm camera field of view for each condition. Embryos were imaged from stage 2-3. Videos were taken simultaneously using two AmScope MD200 USB cameras (AmScope, Irvine, CA) each mounted on an AmScope SE305R stereoscope. Time lapse movies were acquired at a frequency of 1 frame every 10 s for 20 h and saved as Motion JPEG using a MATLAB (The MathWorks, Inc., Natick, MA) script. Movie post-processing (cropping, concatenation, resizing, addition of scale bar) was done using MATLAB and Fiji (Schindelin *et al.*, 2012). All MATLAB scripts written for this study are available upon request. Two of the scripts used here were obtained through the MATLAB Central File Exchange: “videoMultiCrop” and “concatVideo2D” by Nikolay S.

### Embryo nuclei purification

Following *in vitro* fertilization, *X. borealis* embryos were arrested at stage 8 (∼5 hpf) in late interphase using 150 μg/mL cycloheximide in 1/10X MMR for 60 min. Embryos were washed several times in Egg Lysis Buffer (ELB; 250 mM sucrose, 50 mM KCl, 2.5 mM MgCl_2_, and 10 mM HEPES pH 7.8) supplemented with LPC (10 μg/mL each leupeptin, pepstatin, chymostatin), cytochalasin D (100 μg/mL), and cycloheximide (100 μg/mL), packed in a tabletop centrifuge at 200 g for 1 min, crushed with a pestle, and centrifuged at 10,000 g for 10 min at 16°C. The cytoplasmic extract containing endogenous embryonic nuclei was collected, supplemented with 8% glycerol, aliquoted, frozen in liquid nitrogen, and stored at -80°C.

### *X. borealis* egg extract

*X. borealis* metaphase-arrested egg extracts were prepared similarly to *X. laevis* and *X. tropicalis* (Maresca and Heald, 2006; Brown *et al.*, 2007), with the following modifications. *X. borealis* female frogs were primed with 60 U PMSG at least 48 hours before use and boosted with 300 U hCG 12-24 hours before the experiment. Frogs were kept at 16°C in 0.5X MMR. The eggs were dejellied in a 3% cysteine solution in 1X XB-salts (20X: 2M KCl, 20 mM MgCl_2_, 2 mM CaCl_2_), pH 7.8, and washed in CSF-XB (100 mM KCl, 0.1 mM CaCl_2_, 3 mM MgCl_2_, 50 mM sucrose, 10 mM K-EGTA, 10 mM K-HEPES, pH 7.8). CSF-XB buffer was exchanged for CSF-XB supplemented with LPC (10 μg/mL) (CSF-XB+). 1 mL of CSF-XB+ supplemented with cytochalasin D (100 μg/mL) was added to the bottom of an ultracentrifuge tube (SW-55, Beckman) before loading eggs into the tube. Extract tubes were placed in adapter tubes and spun at 1600 g for 1 min to pack the eggs. The excess liquid was removed and the eggs crushed by centrifugation at 10,200 rpm for 16 min at 16°C. The cytoplasmic fraction was then removed and supplemented with LPC (1:1000 dilution; 10 mg/mL stock), cytochalasin D (1:500 dilution; 10 mg/mL stock), energy (1:50 dilution; 50X stock: 190 mM creatine phosphate, 25 mM adenosine triphosphate, 25 mM MgCl_2_, 2.5 mM K-EGTA pH 7.7), and rhodamine-tubulin (1:300 dilution; 65.8 μg/uL stock) for spindle assembly reactions. Sperm nuclei, embryo nuclei, or 10 μm chromatin-coated beads, prepared as previously described (Halpin *et al.*, 2011), were used as a source of DNA for spindle assembly.

### *X. laevis* and *X. tropicalis* egg extracts and mixing experiments

*X. laevis* and *X. tropicalis* metaphase arrested egg extracts were prepared and spindle reactions conducted as previously described (Maresca and Heald, 2006; Brown *et al.*, 2007). *X. laevis* or *X. tropicalis* extracts were mixed in different proportions to *X. borealis* extract and supplemented with either *X. laevis* or *X. tropicalis* sperm nuclei, respectively.

### Extract treatments

To examine sensitivity to perturbations of the RanGTP gradient, *X. laevis*, *X. tropicalis*, and *X. borealis* sperm nuclei were added to their respective egg extracts and cycled through interphase. Ran T24N (final concentration 1 μM) and Q69L (final concentration 5 μM) mutant proteins, purified as previously described (Helmke and Heald, 2014), were added to the interphase extract reaction when fresh CSF extract was added to induce the extract to cycle back to mitosis.

### Spindle spin-down and immunofluorescence

Spindle reactions were prepared, spun-down, and processed for immunofluorescence as previously described (Hannak and Heald, 2006). Briefly, the extract reactions were fixed for 5-10 minutes with 2.5% formaldehyde and spun down at 10,200 rpm for 16 min at 16°C. The coverslips were incubated for 30 s in cold methanol, washed in PBS+NP40, and blocked overnight in PBS + 5% BSA at 4°C. The anti-TPX2 (1:150 dilution, unpublished), anti-katanin p60 (1:500 dilution) (Loughlin *et al.*, 2011), or anti-XMAP215 (1:200, Tony Hyman & Kazu Kinoshita) rabbit antibodies were added for 1 h in PBS + 5% BSA. After washing with PBS+NP40, the coverslips were incubated with 1:1000 anti-rabbit antibody coupled to Alexa Fluor 488 (Invitrogen – Thermo Fisher Scientific, Waltham, MA) for 30 min and then with 1:1000 Hoechst for 5 min. The coverslips were then washed and mounted for imaging with Vectashield (Vector Laboratories, Burlingame, CA).

### Egg whole mount immunofluorescence

Metaphase-arrested *X. borealis* eggs were fixed for 1-3 h at RT using MAD fixative (2 parts of methanol (Thermo Fisher Scientific, Waltham, MA), 2 parts of acetone (Thermo Fisher Scientific, Waltham, MA), 1 part of DMSO), and stored overnight at -20°C in fresh MAD fixative. Embryos were then processed as previously described (Lee *et al.*, 2008) with some modifications. Following gradual rehydration in 0.5X SSC (1X SSC: 150 mM NaCl, 15 mM Na citrate, pH 7.0), embryos were bleached with 2% H_2_O_2_ (Thermo Fisher Scientific, Waltham, MA) in 0.5X SSC containing 5% formamide for 2-3 h under light, then washed in PBT, a PBS solution containing 0.1% Triton X-100 (Thermo Fisher Scientific, Waltham, MA) and 2 mg/ml BSA. Embryos were blocked in PBT supplemented with 10% goat serum (Gibco – Thermo Fisher Scientific, Waltham, MA) and 5% DMSO for 1-3 h and incubated overnight at 4°C in PBT supplemented with 10% goat serum and the primary antibodies. We used the following antibodies: 1:500 mouse anti-beta tubulin (E7; Developmental Studies Hybridoma Bank, Iowa City, IA), 1:350 rabbit anti-histone H3 (ab1791; Abcam, Cambridge, MA). Embryos were then washed 4 X 2 h in PBT and incubated overnight in PBT supplemented with 1:500 goat anti-mouse secondary antibody coupled to Alexa Fluor 488 and 1:500 goat anti-rabbit secondary antibody coupled to Alexa Fluor 568 (Invitrogen – Thermo Fisher Scientific, Waltham, MA). Embryos were then washed 4 X 2 h in PBT and gradually dehydrated in methanol. Embryos were finally cleared in Murray’s clearing medium (2 parts of Benzyl Benzoate, 1 part of Benzyl Alcohol). Embryos were placed in a chamber made using a flat nylon washer (Grainger, Lake Forest, IL) attached with nail polish (Sally Hansen, New York, NY) to a slide and covered by a coverslip, and filled with Murray’s clearing medium for confocal microscopy.

### Confocal microscopy

Confocal microscopy was performed on an upright Zeiss LSM 780 NLO AxioExaminer using the Zeiss Zen software. Embryos were illuminated with an LED light source (X-cite 120LED). For the imaging of histone H3 and tubulin, embryos were imaged using a Plan-Apochromat 20x/1.0 Water objective and laser powers of 12%, on multiple 1024x1024 px plans spaced of 0.683 μm in Z. Images are mean averages of 2 scans with a depth of 16 bits. Channels were acquired simultaneously, and pinhole size was always chosen to correspond to 1 airy unit.

### Imaging and quantification

Eggs were imaged using the Leica Application Suite (v4.9; Leica Microsystems, Buffalo Grove, IL) with a Wild M7A stereoscope equipped with a Leica MC170HD camera. Sperm cells, sperm nuclei, embryo nuclei, and spindles were imaged using micromanager software (Edelstein *et al.*, 2014) with an Olympus BX51 microscope equipped with an ORCA-ER camera (Hamamatsu Photonics, Hamamatsu city, Japan), and with an Olympus UPlan FL 40x/0.75 air objective. All measurements were made using Fiji. The spindle tubulin, katanin, TPX2, and XMAP215 intensity line scans were measured using an automated Java ImageJ plugin developed by Xiao Zhou (Heald lab, UC Berkeley; https://github.com/XiaoMutt/AiSpindle).

### Western blot and analysis

1 μL of egg extract for 3 independent extracts for each species was subjected to SDS-PAGE (12% gel for katanin, 8% gel for TPX2 and XMAP215) and transferred to a nitrocellulose membrane. The anti-TPX2 (1:500 dilution, unpublished), anti-katanin p60 (1:1000 dilution) (Loughlin *et al.*, 2011), or anti-XMAP215 (1:1000, Tony Hyman & Kazu Kinoshita) rabbit antibodies and anti-Ran (1:2000 dilution, BD Biosciences, San Jose, CA) mouse antibody were added for 1 h or overnight at 4°C in PBS + 0.1% Tween-20 + 5% milk. Secondary antibodies (goat anti-rabbit IRDye 800 or goat anti-mouse IRDye 680) were added at 1:10,000 for 1 h in PBS + 0.1% Tween-20. Blots were scanned with an Odyssey Infrared Imaging System (LI-COR Biosciences), and band intensity was quantified with Fiji.

### Protein sequence alignments

Multiple sequence alignments were performed using Clustal Omega (default parameters). Sequence similarities were determined by pairwise alignments using EMBOSS Needle (default parameters).

## ACKNOWLEDGEMENTS

We thank members of the Heald lab for support and fruitful discussions, in particular Xiao Zhou for his line scan plugin and Ambika Nadkarni for the purification of Ran mutants. We thank Sofia Medina-Ruiz, Austin Mudd, and Dan Rokhsar for providing early access to *X. borealis* katanin, TPX2, and XMAP215 sequences. The confocal microscopy performed in this work was done at the UC Berkeley CRL Molecular Imaging Center, supported by NSF DBI-1041078. MK was supported by the UC Berkeley MCB department NIH training grant 4T32GM007232-40. RH was supported by NIH R35 GM118183 and the Flora Lamson Hewlett Chair. RG was supported by an HFSP long term fellowship LT 0004252014-L.

## AUTHOR CONTRIBUTIONS

M.K. performed the experiments and analyzed the data, with help from R.G. M.K. prepared the figures and wrote the manuscript with input from R.H. and R.G.

## AUTHOR INFORMATION

The authors declare no competing financial interests. Correspondence should be addressed to R.G. and R.H.

